# KIMGENS: A novel method to estimate kinship in organisms with mixed haploid diploid genetic systems robust to population structure

**DOI:** 10.1101/2021.10.19.465018

**Authors:** Yen-Wen Wang, Cécile Ané

## Abstract

**Motivation:** Kinship estimation is necessary for evaluating violations of assumptions or testing certain hypotheses in many population genomic studies. However, kinship estimators are usually designed for diploid systems and cannot be used in populations with mixed haploid diploid genetic systems. The only estimators for different ploidies require datasets free of population structure, limiting their usage.

**Results:** We present KIMGENS, an estimator for kinship estimation among individuals of various ploidies, that is robust to population structure. This estimator is based on the popular KING-robust estimator but uses diploid relatives of the individuals of interest as references of heterozygosity and extends its use to haploid-diploid and haploid pairs of individuals. We demonstrate that KIMGENS estimates kinship more accurately than previously developed estimators in simulated panmictic, structured and admixed populations, but has lower accuracy when the individual of interest is inbred. KIMGENS also outperforms other estimators in a honeybee dataset. Therefore, KIMGENS is a valuable addition to a population geneticist’s toolbox.

**Availability and Implementation:** KIMGENS and its association simulation tool are implemented and available open-source at https://github.com/YenWenWang/HapDipKinship.

**Contact:** Yen-Wen Wang

## Introduction

Kinship estimation is crucial to the evaluation of assumption violations (such as when estimating population nucleotide diversity) or to testing various ecological or evolutionary hypotheses (e.g., kin selection). However, kinship estimators for whole genome datasets are mainly developed for human populations (Ramstetter *et al*., 2017). Although these estimators have been widely used in non-human systems, their applications are restricted to diploid-only populations. Nonetheless, a large portion of life forms show plasticity in ploidy (Otto and Gerstein, 2008), which is not accounted for in these estimators. Many plants (e.g. ferns, mosses), fungi (e.g. mushrooms) and algae (e.g. sea lettuces) have complex, multistage life cycles, perform alternation of generations and form haploid structures independent of their diploid counterpart (Brown and Casselton, 2001; John, 1994). Furthermore, most Hymenopterans (e.g. bees and ants), Thysanopterans (e.g. thrips) and some other invertebrates (e.g. some spider mites and rotifers), have an arrhenotokous haplodiploidy system, where males are haploid and females are diploid (Cruickshank and Thomas, 1999; Normark, 2003). Because of the widely present mixed ploidy life-forms, it is crucial to develop estimators that can estimate kinship between individuals with different ploidy levels.

Two marker-based estimators have been developed specifically to estimate relatedness among individuals of different ploidy, including Huang2014, a method-of-moments (MOM) estimator, and Huang2015, a maximum likelihood (ML) estimator (Huang *et al*., 2015). These estimators can thus be used for genome sequencing data directly. In addition, some classical estimators can be extended to estimate relatedness between different ploidies. For example, two kinship estimators, Loiselle 1995 and Ritland 1996 (Loiselle *et al*., 1995; Ritland, 1996), are adapted and implemented in the program PolyRelatedness to resolve inequivalent ploidy (Huang *et al*., 2015). All estimators mentioned above are capable of using multi-allelic loci, which allow them to take advantage of a diversity of genetic markers (e.g., microsatellites). However, these estimators require allele frequencies at each locus in the population, which may not be available in some studies due to sampling strategies (Hahn, 2019). In addition, relying on allele frequencies essentially assumes a population free of stratification. Therefore, the estimators do not account for cryptic population structure, which can result in overestimating in kinship (Manichaikul *et al*., 2010).

To remove the requirement of no population structure, we built on the KING-robust estimator by Manichaikul *et al*., (2010). We extended the estimator’s use to haploid-diploid pairs and named this extension exKING-robust. The exKING-robust estimator uses the heterozygosity of the individuals in the pair of interest as a diversity estimate for background identity-by-descent (IBD). Next, we developed KIMGENS (Kinship Inference for Mixed GENetic Systems), which instead uses the heterozygosity of relatives of the individuals of interest, allowing estimating kinship for diploid, haploid-diploid and haploid pairs of individuals. We showed the estimators are robust to population structure. KIMGENS also performs relatively well under admixture, but can underestimate kinship if an individual is inbred.

## Materials and Methods

We aim to develop a simple kinship estimator that applies to haploid, haploid-diploid and diploid pairs of individuals and is robust to population structure. KING-robust’s strategy is useful for developing a novel kinship estimator for haploid-diploid pairs of individuals. But the strategy cannot apply to haploid pairs directly because it requires the number of heterozygotic sites of an individual to estimate expected heterozygosity (2*pq*) in the ancestral subpopulation of an individual.

To resolve this issue, we propose a two-step approach: (1) We extend KING-robust to obtain a haploid-diploid kinship analysis and to identify a set of diploid relatives for each individual; and (2) for two individuals of interest, *i* and *j*, we use their diploid relatives from step 1 to estimate mean heterozygosity for this pair and to modify their kinship estimate. In the following sections we will first describe a haploid-diploid kinship estimator. Next, we will demonstrate the modification to KING-robust and haploid-diploid kinship estimators for using related individuals *k*. Then, we describe the haploid kinship analysis. Lastly, we will evaluate the performance of these estimators with simulations and a biological dataset (on honeybee).

### Kinship estimation in haploid-diploid pairs of individuals: exKING-robust

The kinship coefficient *ϕ_ij_*, originally termed correlation coefficient of two individuals *i* and *j*, is defined as the probability that two randomly sampled alleles from two individuals are identical-by-descent (IBD) (Lange, 1997; Malécot, 1948). In this section, we derive an estimator for the kinship of a pair of individuals *i_d_* and *j_h_*, where *i_d_* is diploid and *j_h_* is haploid. *ϕ_i_d_j_h__* can be calculated with:

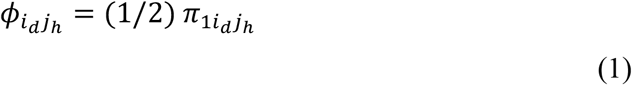

where *π_ni_d_j_h__*denotes the probability of individuals *i_d_* and *j_h_* sharing *n* alleles being IBD. The probability of individual *i_d_* being homozygotic and not in identical-by-state (IBS) with individual *j_h_* at a site can be calculated with:

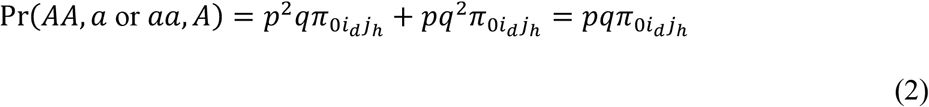

and the probability of individual *i_d_* being heterozygotic and in IBS at the allele in individual *j_h_* at a site can be calculated with:

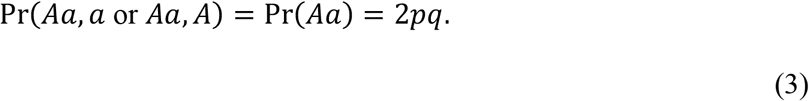

Because *i_d_* and *j_h_* share either 0 or 1 allele by descent (*j_h_* being haploid),

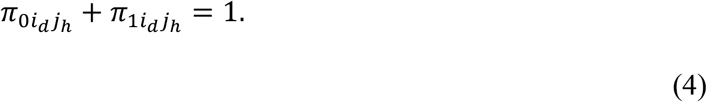

With equation (1), we derive

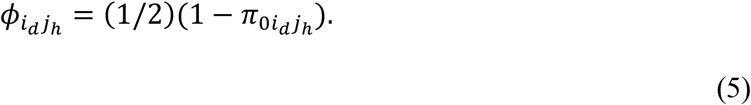

We can combine equation (5) with equation (3) to get

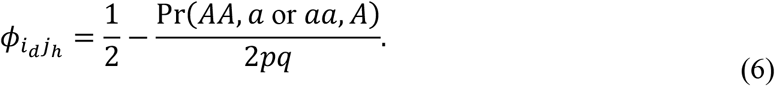

Because only individual *i_d_* is heterozygotic, the expected genome-wide heterozygosity, ∑_*m*_ 2*p_m_q_m_*, can be estimated with 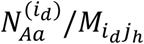 (Manichaikul *et al*., 2010), where 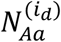 is the number of heterozygotic sites in individual *i_d_* and *M_i_d_j_h__* is the number of sites with non-missing data in both *i_d_* and *j_h_*. ∑_*m*_ Pr(*AA, a* or *aa, A*)_*m*_ can be estimated with *N_AA,a or aa,A_/M_i_d_j_h__*, where *N_AA,a or aa,A_* is the number of sites where individual *i* is homozygotic but not in IBS with individual *j_h_*. Therefore, kinship between individuals *i_d_* and *j_h_* can be estimated with:

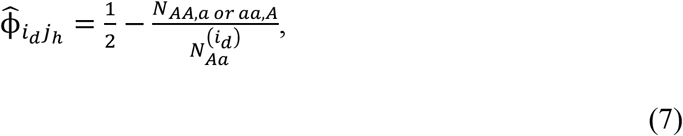

which constitute our exKING-robust estimator for a haploid-diploid pair.

### Methods for using related individuals to estimate *pq*

The KING-robust extension, including KING-robust (Manichaikul *et al*., 2010) for diploid pairs and exKING-robust for haploid-diploid pairs (7), relies on 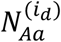 (and 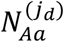). However, in haploid pairs, we do not have the luxury of using the heterozygosity of individuals of interest, so we develop a different estimator, KIMGENS, which uses “heterozygosity references” to estimate *pq*. The accuracy of 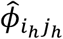 highly depends on the choice of references. To accurately capture heterozygosity, references should come from the same subpopulation as individuals *i_h_* and *j_h_*. Identification of appropriate references can be done by examining kinship estimates between the individual *i_h_* and *j_h_* and the potential references. Since some individuals may deviate from Hardy-Weinberg equilibrium (HWE) in subpopulations, choosing a single reference from the relatives of either individual *i_h_* or *j_h_* may result in using an inbred or admixed product, biasing the estimate. So, we choose two sets of individuals *K*(*i_h_,t*) and *K*(*j_h_,t*), which are related to either one of the two individuals of interest, given a kinship threshold *t*. Every individual *k* in *K*(*i_h_,t*) or *K*(*j_h_,t*) is used as a heterozygosity reference for an intermediate kinship estimate, 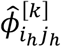. Then, we calculate two medians of intermediate kinship estimates, one from *K*(*i_h_,t*) and one from *K*(*j_h_,t*). Finally, we take the mean of these two medians as our final estimate 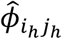. We explain this estimation procedure below in detail.

For more generality, we introduce this procedure for diploid individuals to modify the exKING-robust estimators as well. For two diploids *i_d_* and *j_d_* and for a reference individual *k*, we define the intermediate kinship estimate 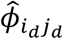 as

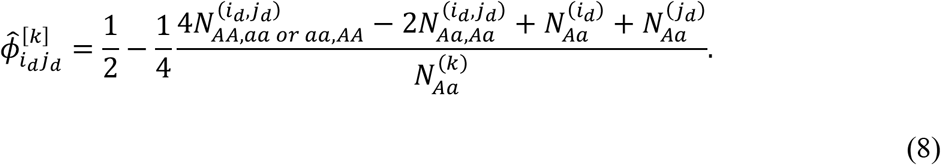

Next, for an individual *x*, we consider its references to be the set *K*(*x,t*) of diploid individuals that share kinship with *x* greater than a given threshold *t* (including *x* itself if *x* is diploid), based on the exKING-robust kinship estimate. Finally, we define the KIMGENS estimate as follows:

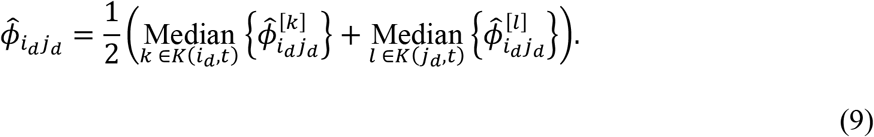

Parenthesized superscripts denote the individuals with which sequences are compared to derive the number of sites with a particular pattern, and bracketed superscripts denote the individuals used for intermediate kinship estimates. Note, (8) corresponds to equation (11) in Manichaikul *et al*., (2010) for *k* taken to be either *i_d_* or *j_d_*, whichever has the smallest 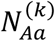. The innovation here is to consider the median of kinship estimates, and to use close relatives (not just *i_d_* or *j_d_*) to approximate heterozygosity at the denominator.

Using the same idea, we use (7) to define the intermediate kinship estimate between a pair of diploid and haploid individuals, given a reference individual *k* as:

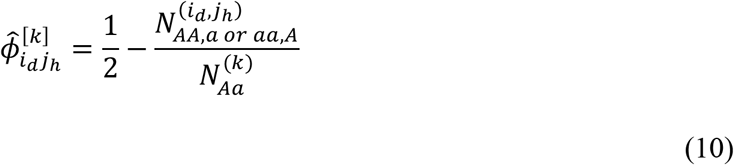

and for a haploid-diploid pair we define the KIMGENS estimate as:

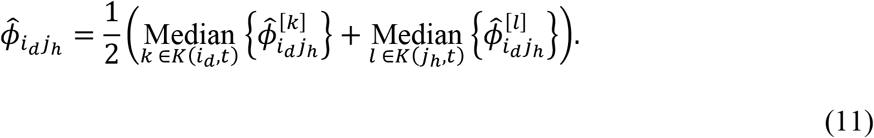

When calculating a 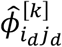 (or 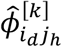), there are three individuals involved: *i_d_*, *j_d_* (or *j_h_*) and *k*. The amount of missing data are not the same in these three individuals. So, we only consider the sites that are non-missing in all three individuals for each 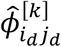 (or 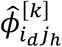).

### Kinship estimation in haploid pairs of individuals

Under the same definition for kinship, in haploid pairs, the kinship coefficient *ϕ_i_h_j_h__* can be calculated with:

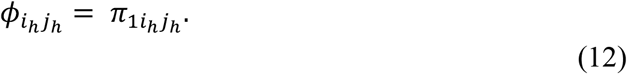

The probability of individuals *i_h_* and *j_h_* not in IBS at a site can be calculated with:

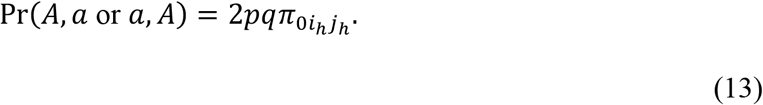

Because

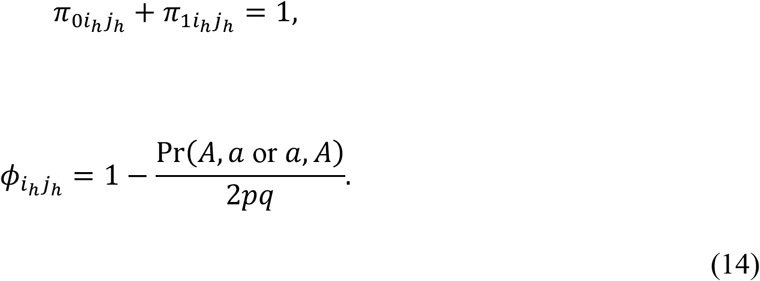

Using the same strategy described above, an intermediate kinship for haploid pairs of individuals can be estimated using a reference diploid individual *k* with:

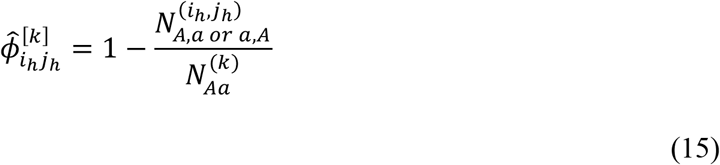

and the KIMGENS estimate for a haploid-haploid pair is defined as

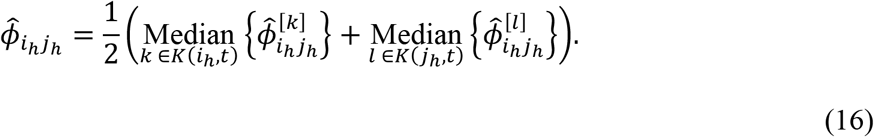

### Simulations

To assess the performance of these estimators, we simulated panmictic, structured and admixed populations of species with haplodiploid or diploid genetic system. For panmictic populations, the allele frequency of each site was simulated from a uniform distribution between 0.1 and 0.9, U(0.1,0.9). The genotypes for starter individuals (those without known parents) in pedigree simulations were drawn from the allele frequency. For structured and admixture populations, the allele frequencies of three subpopulations were simulated following the Balding-Nichols model from a panmictic ancestral population (allele frequency drawn from U(0.1,0.9)). The Wright’s Fst (θ_k_) of the subpopulations was set to 0.05, 0.15 and 0.25. In structured populations, each family was drawn from a random subpopulation. To simulate admixture, Conomos’s strategy was used (Conomos *et al*., 2016). In pedigree simulations, the ancestry proportions of the founders were drawn independently from either of two Dirichlet distributions: Dir(6, 2, 0.3) and Dir(2, 6, 0.3), and the genotypes of the founders were drawn from the ancestry and allele frequencies of the three subpopulations.

While simulating pedigrees, nine different scenarios were simulated 1000 times each.

The scenarios differed by four factors: (1) the number of independent SNP sites: 20k or 100k, (2) genetic system: arrhenotokous haplodiploidy or diploidy, (3) population structure: panmictic, structured or admixture and (4) pedigrees (Supplementary Table 1). Overall, 100k SNP sites were simulated unless when the estimators being compared included those implemented in PolyRelatedness, in whick case 20k SNPs were simulated. All simulations are under haplodiploidy unless otherwise noted. First, to evaluate the performance of exKING-robust and KIMGENS, we simulated a single large family (Supplementary Figure 1) from a panmictic population (scenario 1). To compare the performance with that of previously published estimators, we simulated 11 families (Supplementary Figure 2) from a panmictic or structured population (scenarios 2 and 3). To explore the performance of the estimators under admixture, we simulated the single large family (Supplementary Figure 1) or 11 families (Supplementary Figure 2) from an admixed population with 100k or 20k sites (scenarios 4 and 5). To understand how different estimators perform on inbred products, we simulated five inbreeding families (Supplementary Figure 3) and ten unrelated individuals (so PolyRelatedness can estimate allele frequency more accurately) from a panmictic diploid or haplodiploid population (scenarios 6 and 7). Lastly, to explore the use of different thresholds (*t*), we simulated a new family (Supplementary Figure 4) with twenty unrelated diploid individuals in a structured or admixed population (scenarios 8 and 9).

We estimated pairwise kinships for all individuals using different estimators and extracted the estimates of the pairs of interest. To convert relatedness (calculated by the estimators implemented in PolyRelatedness) to kinship, the relatedness estimates for diploid pairs were divided by two and those for haploid-haploid and haploid-diploid pairs were divided by one. To summarize the estimation, we calculated the bias 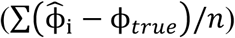 and root-mean-square error (RMSE; 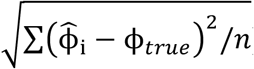) of each estimator. For KIMGENS, the threshold *t* was arbitrarily set to 0.1, except when exploring different thresholds, in which case *t* was set to either 0.1 or 0. Inbreeding coefficients were calculated from pedigrees with the R package kinship2 (Sinnwell *et al*., 2014).

### Biological data

In addition to simulations, we used a honeybee dataset which was originally collected for estimating crossover rate (Liu *et al*., 2015). This dataset includes three monogynous colonies (one queen per colony). One queen (diploid) and multiple drones (haploids) were sampled from all three colonies. Six additional workers (diploids) were sampled from one of the colonies. Also, three drones were sequenced twice. We therefore expect that from a single colony, (1) the drones and the queen share a kinship of 0.5, (2) the workers and the queen share a kinship of 0.25, (3) the drones share a kinship of 0.5 with each other, (3) the workers share a kinship of 0.375 (full-siblings) or 0.125 (half-siblings) with each other, (4) the drones and workers share a kinship of 0.25, and (5) the two sequences from the same drone share a kinship of 1 with each other.

The genomic raw reads were downloaded from NCBI and mapped to reference genome (GCF_000002195.4) with BWA mem ver. 0.7.17. Duplicated reads were filtered with samtools ver. 1.9 and SNPs were called with bcftools ver. 1.9. To avoid identifying SNPs due to indels, we applied four filters: (1) the repetitive regions identified by RepeatMasker, (2) sites with read depth higher than 1.3X mean depth or lower than 0.75X mean depth, (3) sites with minor allele frequency lower than 0.01 and (4) sites that are called heterozygous in any haploid individuals (drones). All filtered SNPs (N=1,008,683) were used to estimate kinship without LD correction. As the previous section, for KIMGENS, the threshold *t* was arbitrarily set to 0.1. To compare KIMGENS with other published estimators, we sampled one every twenty SNPs to avoid segmentation faults for these other estimators.

## Results and discussion

### Evaluation of the methods under a panmictic population

We simulated a single large family in haplodiploidy with 100k sites in a panmictic population (Supplementary Table 1 and Supplementary Figure 1) and compared the performance of exKING-robust and KIMGENS, for each ploidy level of the individuals of interests (diploid, haploid-diploid or haploid). For diploid pairs of individuals, the estimates from both exKING-robust and KIMGENS are accurate with no bias and small RMSE (Figure 1, Supplementary Table 2). The same is observed for haploid-diploid and haploid pairs (Figure 1, Supplementary Table 2). The variance of estimates is usually higher in haploid pairs and lower in diploid pairs, likely due to the fact that the amount of allelic data is halved in haploid compared to diploid individuals, causing a precision decrease.

**Figure 1.**
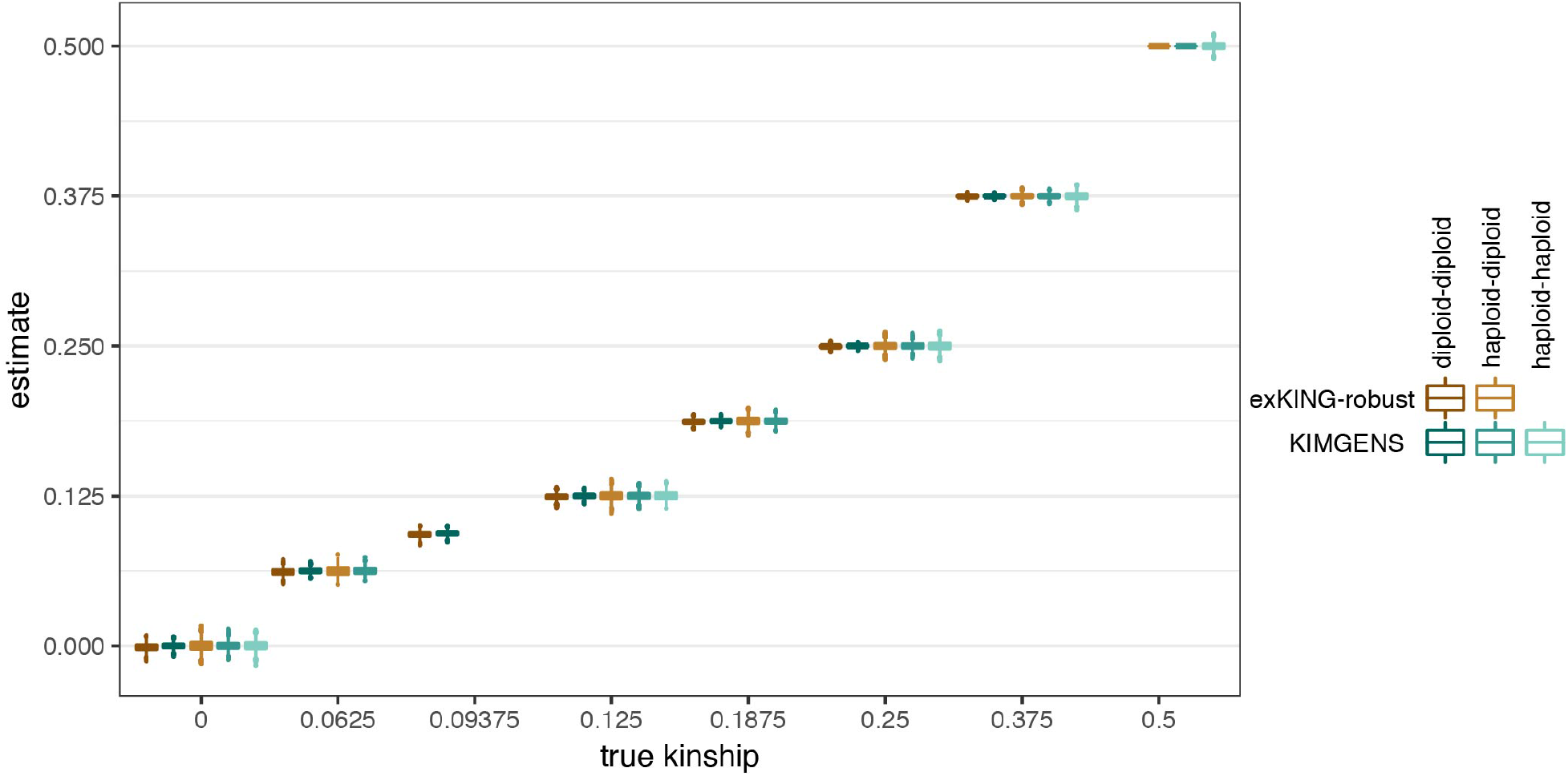
Distribution of kinship estimates of exKING-robust and KIMGENS in a panmictic population. Boxplots show the median, first and third quartiles, and range of each distribution.

### Comparison with previous methods in panmictic and structured populations

We compared the performance of KIMGENS with other relatedness estimators implemented in the package PolyRelatedness, including Huang2014 (MOM), Huang2015 (MLE), Ritland 1996 and Loiselle 1995 (Huang *et al*., 2015; Loiselle *et al*., 1995; Ritland, 1996). We simulated 11 families from a panmictic or structured population (Supplementary Table 1 and Supplementary Figure 2).

In a panmictic population, the performance of KIMGENS outcompetes all other estimators in terms of the overall RMSE and bias (Figure 2A, Supplementary Table 3). KIMGENS performs slightly worse than Huang 2015 only when the true kinship is zero. In a structured population, KIMGENS again outperforms all other methods when the true kinship is not zero (Figure 2B and Supplementary Table 4). However, KIMGENS has the highest RMSE and absolute bias when true kinship is zero, and Huang 2015 has the lowest. In the structured population simulation, there is 1/3 chance that two unrelated individuals are from different subpopulations. The fixed variants in the subpopulations increase the homozygotic differences between two individuals and hence lower the kinship estimates between unrelated samples using KING-robust-based strategies (Manichaikul *et al*., 2010). Also, note that Huang 2015 performs the best when the true kinship equals zero in both conditions (Supplementary Table 4). This is likely because Huang 2015 uses a maximum likelihood strategy searching for IBD on a parameter space, where the lower bound of the parameter space is zero (Huang *et al*., 2015). If negative kinship is a concern, one can enforce a lower bound of zero for all estimators. In our simulations, this would vastly improve the bias and RMSE of KIMGENS when the true kinship is zero, without affecting the performance when the true kinship is positive.

**Figure 2.**
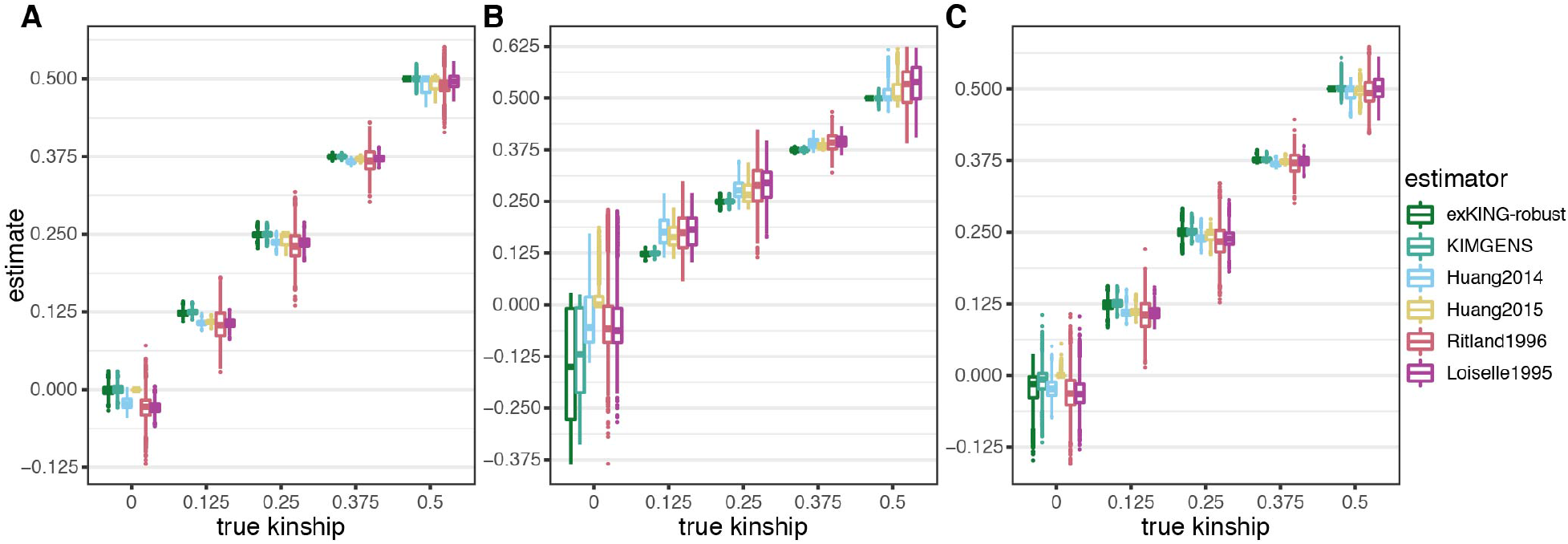
Kinship estimates of different estimators in panmictic (A), structured (B) and admixed (C) populations.

### Estimates in an admixed population

Although estimating kinship in an admixed population is not the goal of this project, we explored the robustness of KIMGENS in admixed population. First, we simulated the single-family pedigree in an admixed population to evaluate the performance of exKING-robust and KIMGENS (Supplementary Table 1 and Supplementary Figure 1). Like previous reports on KING-robust (Conomos *et al*., 2016), the accuracy of both estimators drops compared to the estimates in a panmictic population because the individuals from a single family may have different ancestries (Supplementary Table 5 and Supplementary Figure 5). Similarly to the panmictic population simulation, the estimates in haploid and diploid pairs of individuals have slightly higher and lower RMSE, respectively. KIMGENS also performs slightly better than exKING-robust in terms of RMSE and bias.

We further compared KIMGENS with aforementioned estimators using the 11-family-pedigree (Supplementary Table 1 and Supplementary Figure 2). In this admixture simulation, KIMGENS has lower absolute bias than all other estimators but a RMSE slightly higher than Huang 2015 (Figure 2C and Supplementary Table 6). The relatively high RMSE is also driven by the lower kinship estimates on unrelated individuals due to the same reasons discussed in the last section.

### Estimates on inbred individuals

Like KING-robust, KIMGENS is not designed to calculate kinship in inbred populations, but we explored its performance for inbred individuals by simulating five families and ten additional unrelated individuals (half male and half female) in a panmictic population in diploid or haplodiploid genetic system (Supplementary Table 1 and Supplementary Figure 3). The unrelated individuals were included because all of the methods being compared require population allele frequency, which is estimated with the sampled individuals in this study. In a diploid genetic system, KIMGENS performs slightly better than other estimators overall in terms of both RMSE and bias (Figure 3A and Supplementary Table 7). However, the RMSE and absolute bias increase when the individual inbreeding coefficients of the two individuals increase, and the increasing rate is faster than other kinship estimators, such as Huang 2015 and Loiselle 1995.

**Figure 3.**
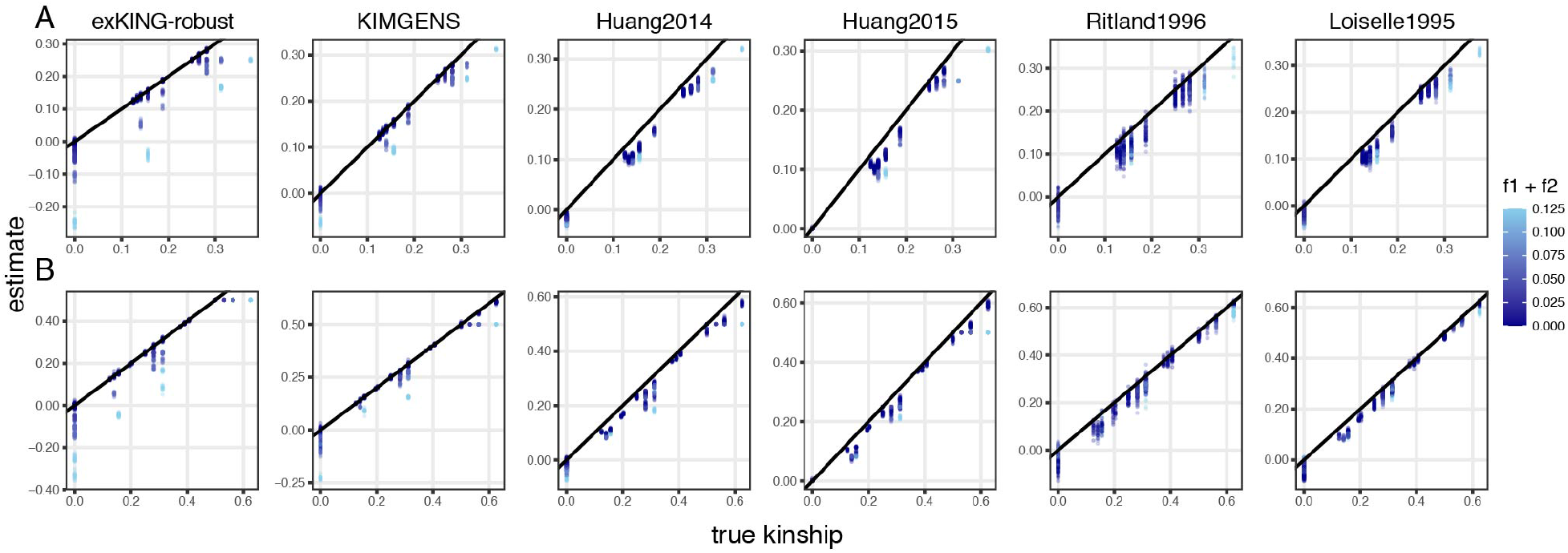
Performance of different estimators on inbred diploid (A) and haplodiploid (B) populations. A thousand points were chosen randomly to be presented on each plots. Diagonal line: estimated kinship = true kinship. f1+f2: the sum of inbreeding coefficients of the two individuals in a pair of interest.

In a haplodiploid genetic system, KIMGENS has a relatively high overall RMSE and absolute bias (Figure 3B and Supplementary Table 8), so we broke down the results by the ploidy of pairs and individual inbreeding coefficients. For diploid pairs, the behavior of KIMGENS is very similar to that in the diploid simulation (Supplementary Table 8 and Supplementary Figure 6A). KIMGENS outperforms all other estimators overall, but the accuracy drops when the individual inbreeding coefficient increases. For haploid pairs, all individuals have zero inbreeding coefficients and KIMGENS also performs better than other estimators (Supplementary Table 8 and Supplementary Figure 6C). However, for haploid-diploid pairs, KIMGENS performs worse than other estimators overall except for exKING-robust and also when individual inbreeding coefficients are higher than zero (Supplementary Table 8 and Supplementary Figure 6B). Like diploid pairs, the kinship estimates decrease under a higher degree of inbreeding (Supplementary Table 8 and Supplementary Figure 6B). This correlation is essentially the same underestimation as when the individuals in the pair of interest are from two different subpopulations.

### Performance on biological data

The kinship estimates on the honeybee dataset using KIMGENS are close to expectations in all within-colony relationships except for workers-workers (Figure 4; Supplementary Table 9). Kinship estimates between workers can be clustered into three groups: 0.125, 0.25 and 0.375. While estimates at 0.125 and 0.375 between workers are expected for full-siblings and half-siblings, estimates at 0.25 are unexpected, but is likely the result of paternal relatedness. For example, the kinship between two workers whose fathers are siblings equals 0.25. In addition, we found that individuals between different colonies share a considerably high degree of kinship (mean= 0.08) (Figure 4). We hypothesize that the high degree of kinship is derived from true background relatedness due to breeding management of the bee farm. The background relatedness may also contribute to the positive biases of kinship estimates between workers within a single family-that is, the putative half-siblings may have distantly related fathers (Supplementary Table 9).

**Figure 4.**
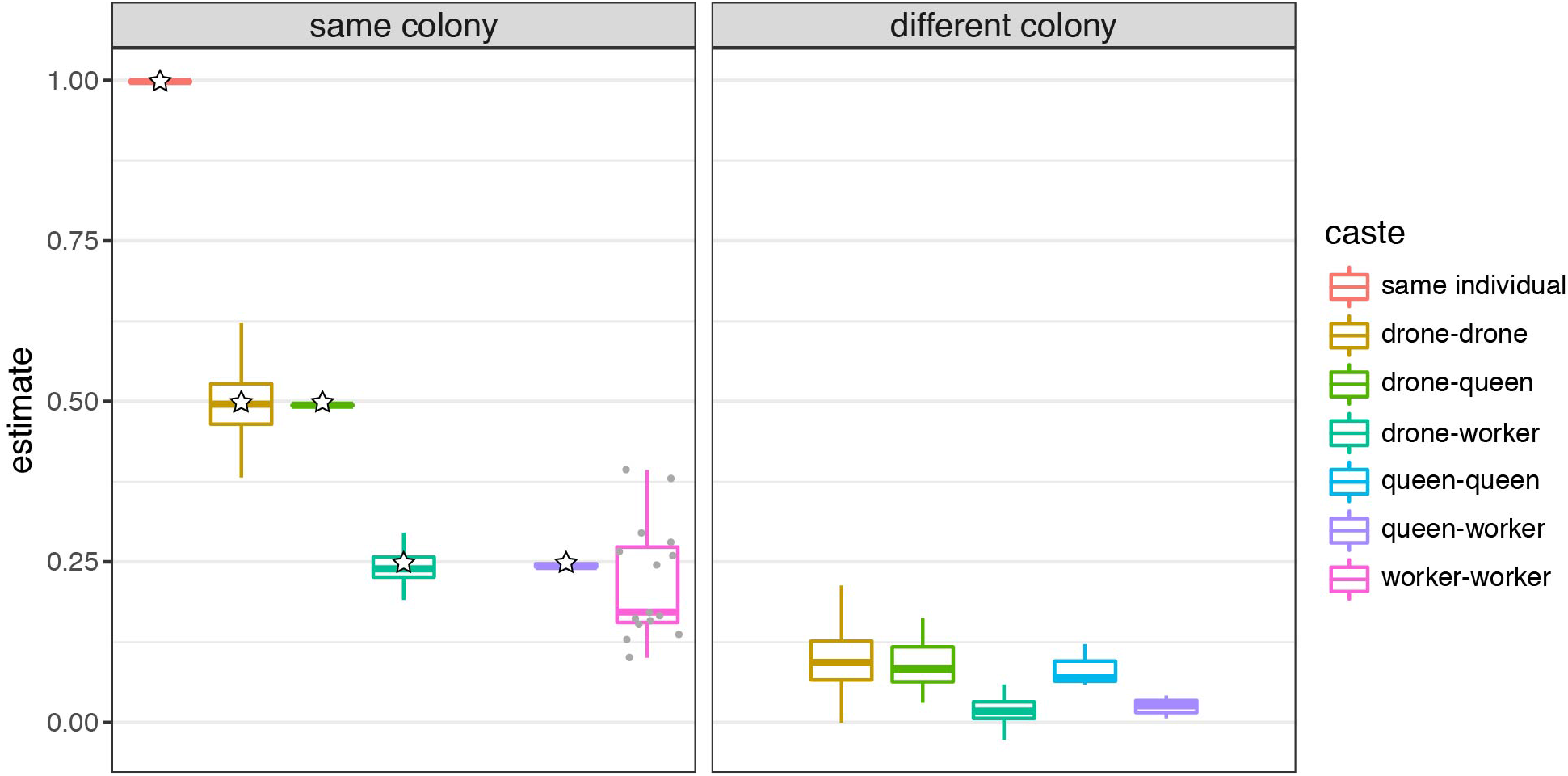
Kinship estimates from KIMGENS on three honeybee colonies. Gray dots indicate each kinship estimate between workers. The “same individual” category consists of pairs of sequence data sets from the drones that were sequenced twice. Stars indicate expected kinship.

The performance of KIMGENS on the subsampled dataset is similar to that on the whole dataset, while all other estimators underestimate kinships on the subsampled dataset when the true kinships are lower than 1 (Supplementary Figure 7 and Supplementary Table 10). This observation supports the usage of KIMGENS on biological datasets.

### Choice of the kinship threshold *t*

The only parameter in KIMGENS is the kinship threshold *t* used to define the set of relatives for heterozygosity referencing. Without inbreeding in a panmictic or structured population, the threshold *t* should not affect the accuracy as long as it is positive. However, admixture may elevate unrelated individuals kinship (Figure 2C), and inbreeding can lower the heterozygosity in some individuals, so the choice of *t* needs to be taken into consideration.

The threshold *t* can affect two factors: the accuracy of heterozygosity referencing and the number of reference individuals. In order to identify diploid individuals that can represent the heterozygosity of the individuals of interest (in a same population), one should consider a higher *t*, but a higher *t* may reduce the number of heterozygosity references. A lower number of references should not directly affect the accuracy of kinship estimation. For example, although there are 46 drones in the honeybee dataset and only nine females, the female honeybees are closely related to the drones and hence can represent the heterozygosity of the drones well. However, if a dataset includes numerous inbreeding events, using a high *t* may result in only referencing the inbred individuals, and the kinship estimation will be inaccurate, so a lower *t* should be considered. In some extreme cases, it may be that KIMGENS cannot estimate kinship, if zero diploid relatives are available for a pair of haploid individuals. In this case, one may choose zero for threshold *t*, which allows referencing unrelated individuals from the same subpopulation for heterozygosity.

To explore the effect of using unrelated individuals as heterozygosity references on kinship estimation for haploid pairs of individuals, we simulated one family and twenty unrelated diploids in a structured or admixed population (Supplementary Table 1 and Supplementary Figure 4). We first estimated kinships regularly using a threshold *t* at 0.1 with KIMGENS. Then, we removed all diploid individuals within the family, leaving the unrelated diploids in the dataset only and estimated kinship using a threshold *t* at 0. In a structured population, both the RMSE and absolute bias are higher when using non-relatives compared to using relatives, but the estimates are still accurate (Supplementary Table 11 and Supplementary Figure 8A). In admixed populations, the RMSE and bias are also higher when using non-relatives; however, the RMSE is noticeably higher when the true kinship is over zero (Supplementary Table 11 and Supplementary Figure 8B). This is likely due to the complex ancestry of the individuals in admixed populations, resulting in non-relatives providing a significantly worse heterozygosity reference than relatives. Of note, in 1% of the pairs of interest, there were no diploid individuals with a kinship above 0 (but less than 0.1) with either haploid individual in the pair, so no kinship was estimated for these pairs. Using a negative threshold *t* can resolve the issue, but these estimates should be interpreted with extra caution.

## Conclusions

Here we present new kinship estimators for mixed haploid-diploid populations that are robust to population structure. We demonstrate the accuracy of KIMGENS in panmictic, structured and admixed simulated populations as well as in a biological dataset. Simulations and biological datasets indicate that KIMGENS performs better than previously developed kinship estimators, but one may choose to use previously developed kinship estimators when the dataset contains many multiallelic loci or individuals of interest with high degree of inbreeding coefficient. The methods are implemented in an R package available on github (https://github.com/YenWenWang/HapDipKinship) for researchers studying population genomics in mixed ploidy systems.

## Data Availability Statement

The data underlying this article are available in Open Science Repository at https://dx.doi.org/10.17605/OSF.IO/EP6MF. The datasets were derived from NCBI sequence read archive, accession SRP043350.

## Acknowledgements

The authors thank Dr. Anne Pringle and Dr. Jacob Golan for reviewing the manuscript, and Dr. Cameron Currie and Dr. Sean Schoville for advice on biological datasets.

## References

Brown, A.J. and Casselton, L.A. (2001) Mating in mushrooms: Increasing the chances but prolonging the affair. Trends Genet., 17, 393–400.

Conomos, M.P. et al. (2016) Model-free estimation of recent genetic relatedness. Am. J. Hum. Genet., 98, 127–148.

Cruickshank, R.H. and Thomas, R.H. (1999) Evolution of haplodiploidy in dermanyssine mites (Acari: Mesostigmata). Evolution (N. Y)., 53, 1796–1803.

Hahn, M. (2019) Experimental design. In, Molecular population genetics. Sinauer Associates, Inc., Sunderland, Massachusetts, pp. 25–42.

Huang, K. et al. (2015) Estimating pairwise relatedness between individuals with different levels of ploidy. Mol. Ecol. Resour., 15, 772–784.

John, D.M. (1994) Alternation of generations in algae: Its complexity, maintenance and evolution. Biol. Rev. Camb. Philos. Soc., 69, 275–291.

Lange, K. (1997) Mathematical and statistical methods for genetic analysis. Springer, New York, NY, USA.

Liu, H. et al. (2015) Causes and consequences of crossing-over evidenced via a high-resolution recombinational landscape of the honey bee. Genome Biol., 16, 15.

Loiselle, B.A. et al. (1995) Spatial genetic structure of a tropical understory shrub, Psychotria officinalis (Rubiaceae). Am. J. Bot., 82, 1420–1425.

Malécot, G. (1948) Les mathématiques de l’hérédité. Masson et Cie, Paris, France.

Manichaikul, A. et al. (2010) Robust relationship inference in genome-wide association studies. Bioinformatics, 26, 2867–2873.

Normark, B.B. (2003) The evolution of alternative genetic systems in insects. Annu. Rev. Entomol., 8, 397–423.

Otto, S.P. and Gerstein, A.C. (2008) The evolution of haploidy and diploidy. Curr. Biol., 18, R1121–R1124.

Ramstetter, M.D. et al. (2017) Benchmarking relatedness inference methods with genome-wide data from thousands of relatives. Genetics, 207, 75–82.

Ritland, K. (1996) Estimators for pairwise relatedness and individual inbreeding coefficients. Genet. Res., 67, 175–185.

Sinnwell, J.P. et al. (2014) The kinship2 R package for pedigree data. Hum. Hered., 78, 91–93.

